# Circulating MicroRNAs as Potential Diagnostic Biomarkers for Heart Failure: A Systematic Review and Meta-Analysis

**DOI:** 10.64898/2026.03.09.710696

**Authors:** Wenqing Sun, Bangqin Hu, Daiyi Li, Yan Qian, Chengzhi Huang, Na Wang

**Affiliations:** Department of Pharmacy, The Second Affiliated Hospital of Chongqing Medical University, Chongqing 400010, P. R. China; Key Laboratory of Luminescence Analysis and Molecular Sensing (Southwest University), Ministry of Education, College of Pharmaceutical Sciences, Southwest University, Chongqing 400715, P. R. China; Department of Pharmacy, Department of Pharmacy, The First Affiliated Hospital of Jinan University,GuangZhou, 510000, PR China

**Keywords:** Meta-Analysis, Heart Failure / diagnosis, MicroRNAs / blood, Biomarkers, Systematic Review

## Abstract

**Background:** Current heart failure (HF) biomarkers (e.g., BNP/NT-proBNP) have limitations in specificity and performance in HF with preserved ejection fraction (HFpEF). Circulating microRNAs (miRNAs) are promising novel biomarkers. This study aimed to comprehensively evaluate the diagnostic stability of circulating miRNAs for HF, identify novel candidates, and prioritize them for clinical translation.

**Methods:** We conducted a systematic review and meta-analysis. PubMed, Embase, and Cochrane Central were searched from inception to March 2025. Studies comparing miRNA expression in HF versus control groups using blood or tissue samples were included. Data were extracted, and study quality was assessed using the Newcastle-Ottawa Scale (human) and SYRCLE’s tool (animal). A random-effects model pooled log odds ratios (logORs) for each miRNA. Subgroup analyses were based on species, ethnicity, and sample type. Evidence quality was graded using the GRADE framework.

**Results:** Eighty-six studies (61 human, 25 animal) with 3,023 samples were included. Meta-analysis identified 71 consistently dysregulated circulating miRNAs (58 up, 13 down) in HF. Key upregulated miRNAs included miR-21 (logOR=8.15, 95% CI: 7.55–8.74), miR-423-5p, and miR-210. Key downregulated miRNAs included miR-144 and miR-126. Subgroup analyses revealed differences by species, ethnicity (Asian vs. non-Asian), and sample type (serum vs. plasma). GRADE assessment classified five miRNAs (miR-1, miR-21, miR-221, miR-423-5p, miR-148a) as high-quality evidence.

**Conclusions:** This meta-analysis identifies a panel of circulating miRNAs with stable expression in HF, with miR-21 and miR-423-5p being the most robust. Evidence grading provides a clear priority list (e.g., miR-21, miR-423-5p) for clinical validation. Subgroup heterogeneity highlights the need for standardized protocols and precision diagnostics in future biomarker development.

## Introduction

Heart failure (HF), the end stage of numerous cardiovascular diseases, remains a leading cause of high morbidity and mortality worldwide^1,2^. Despite advances in diagnosis and treatment, established biomarkers such as B-type Natriuretic Peptide (BNP) and N-terminal pro-B-type Natriuretic Peptide (NT-proBNP) have important limitations.their levels are influenced by confounding factors including age, renal impairment, obesity, and atrial arrhythmias^3,4^ and their diagnostic accuracy in HF with preserved ejection fraction (HFpEF) remains uncertain^5^. These constraints highlight the need for novel biomarkers with greater specificity and robustness, particularly for early detection.

In recent years, microRNAs (miRNAs) have emerged as potential diagnostic biomarkers and therapeutic targets in cardiovascular disease, including HF. As key post-transcriptional regulators, circulating miRNAs modulate critical pathological processes such as myocardial hypertrophy, fibrosis, apoptosis, and ventricular remodeling^6–8^. Their relative stability in peripheral blood and tissue-specific expression profiles make them attractive candidates for noninvasive diagnostis^9–11^ and risk stratification in HF^2,12,13^. Numerous studies have reported characteristic alterations in the circulating miRNA expression profile in HF patients. For instance, miRNAs such as miR-21, miR-23a, miR-142-5p, miR-126, and miR-195 have been associated with myocardial injury and deterioration of cardiac function^9,14–16^.

Since the publication of earlier systematic reviews^9,17^, substantial new evidence has emerged, with at least 31 additional high-quality studies now available^10,14,18–47^. Previous meta-analyses were limited by the number of included studies, preventing comprehensive subgroup analyses on crucial factors like ethnicity and sample type. Recent studies ^14,48^ now permits more comprehensive evaluation of these variables.

This systematic review and meta-analysis aimed to: 1) assess the diagnostic stability of previously identified core miRNAs in a larger dataset; 2) identify novel candidate miRNAs and elucidate their potential application scenarios in specific populations via subgroup analysis; and 3) screen for high-priority miRNA molecules with greater potential for clinical application. These findings are expected to establish a refined, high-priority list of miRNA candidates for the early diagnosis of heart failure and facilitate their translation into clinical practice.

## Methods

### Literature Search and Selection

To comprehensively identify relevant studies, two researchers independently conducted a systematic search in the PubMed, Embase, and Cochrane Central Register of Controlled Trials for literature published from the inception of each database until March 25, 2025, limiting the search to English publications. The search strategy combined subject headings with free-text words, with core concepts including (“microRNA” OR “miRNA” OR “miR-”), (“heart failure” OR “HF”), and (“expression” OR “analysis” OR “profile”). The complete search strategy is provided in the Supplementary Materials. After initial screening, the search results were cross-checked by the two reviewers. Any discrepancies were resolved through discussion with a third researcher until a consensus was reached.

### Inclusion and Exclusion Criteria

Studies were eligible for inclusion if they met all of the following criteria: (1) original studies comparing miRNA expression levels between a heart failure group and a healthy (or non-heart failure) control group; (2) miRNAs derived from blood samples (plasma, serum, or whole blood) or myocardial tissue; (3) provision of extractable miRNA expression data and sufficient sample size information; (4) miRNA detection using established techniques such as quantitative PCR, real-time quantitative PCR, or microarray; and (5) published as a full-text article in English.

Studies were excluded if any of the following applied: (1) study population complicated by other severe diseases (e.g., malignant tumors, end-stage renal disease); (2) purely basic research (e.g., involving only cell experiments); (3) non-original research types, including letters, commentaries, case reports, meta-analyses, editorials, etc.; (4) data missing or key information unavailable.

### Data Extraction and Quality Assessment

Data were extracted from the included studies using a pre-designed standardized form. Extracted information included: first author, year of publication, ethnicity of the study population, species, miRNA detection method, specific sample type, sample size, and the names and direction of expression of differentially expressed miRNAs.

Methodological quality assessment was performed using different tools according to the type of study objects:

For human studies, the Newcastle-Ottawa Scale (NOS)^49^ was employed. This scale evaluates study quality across three domains—selection of study populations, comparability between groups, and assessment of exposure/outcome—through a total of eight items, using a star-based system (maximum 9 stars). Two independent researchers conducted the ratings, and any discrepancies were resolved through discussion or consultation with a third researcher.

For animal experimental studies, the SYRCLE’s risk of bias tool was applied^50^. This tool is an adaptation of the Cochrane risk of bias tool tailored to the characteristics of animal experiments. It assesses the internal validity of studies across six domains—selection bias, performance bias, detection bias, attrition bias, reporting bias, and other biases—comprising 10 specific items. Each item was judged as “yes” (low risk), “no” (high risk), or “unclear” based on the information reported. The assessment was carried out independently by two researchers, and disagreements were resolved through discussion or by consulting a third researcher until consensus was reached.

### Data Synthesis and Statistical Analysis

All statistical analyses were performed using STATA 16. A random-effects model was employed to pool effect sizes. The logarithmic odds ratio (logOR) with corresponding 95% confidence interval (CI) were used as the summary statistic to evaluate the expression difference of each miRNA between the heart failure and control groups. Positive logOR values with 95% CI excluding zero indicated significant upregulation in HF, whereas negative values indicated downregulation. Statistical significance was defined as P < 0.05. For the identified differentially expressed miRNAs, we comprehensively evaluated their potential importance based on the following three dimensions: (1) the number of studies reporting significant differences for the miRNA; (2) the total sample size supporting the conclusion; and (3) the absolute magnitude and statistical significance of the pooled logOR value. Furthermore, subgroup analyses were performed based on species (human/animal), sample type (blood sample/tissue sample), and ethnicity (Asian/non-Asian). Sensitivity analyses were conducted to assess the robustness of the findings and sources of heterogeneity.

### Evidence Quality Grading

The quality of evidence for each differentially expressed miRNA was graded using the GRADE (Grading of Recommendations Assessment, Development, and Evaluation) system^51^.

The initial evidence level was determined based on study design: human observational studies started as “low” quality, whereas animal studies began as “very low” quality due to inherent indirectness between animal models and humans.

The grading process involved evaluating five predefined domains. Factors that could lower the evidence quality included: risk of bias (if ≥50% of the included studies had a high risk of bias), inconsistency (substantial unexplained heterogeneity across studies, I² > 50%), indirectness (mismatch in population, samples, or assay methods relative to clinical application scenarios), imprecision (wide 95% confidence interval of the pooled effect size or total sample size < 100 cases), and publication bias. Conversely, evidence could be upgraded in the presence of a large effect size (absolute value of pooled lgOR > 9.00 with lower 95% CI limit > 7.00), a potential dose-response relationship, or evidence suggesting that residual confounding would underestimate the true effect.

Finally, the body of evidence for each miRNA was classified into one of four quality levels: high, moderate, low, or very low. A high grade indicates high confidence in the effect estimate, whereas a very low grade reflects considerable uncertainty. All assessments were performed independently by two researchers, and any discrepancies were resolved through consensus.

## Results

### Literature Search and Basic Characteristics of Included Studies

The initial search identified 3702 potentially relevant articles. After removing duplicates and initial screening, 102 articles underwent full-text review. Ultimately, 86 articles (comprising 3023 total samples) were included in the systematic review and meta-analysis. The detailed literature screening process is illustrated in Figure 1.

**Figure 1.**
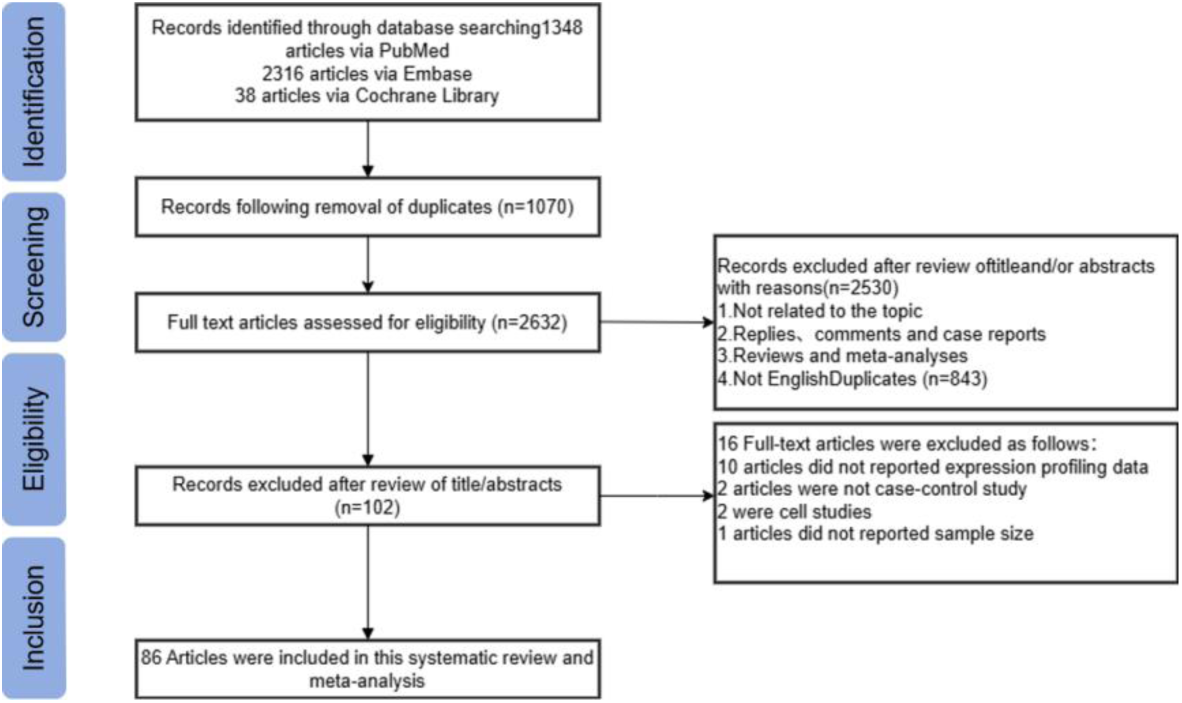
Flow diagram of the study selection process.

The included studies consisted of 61 human studies (2812 samples) and 25 animal studies (211 samples). The animal studies involved rats, mice, sheep, and dogs. The human studies encompassed populations from multiple global regions, including China, Italy, the United States, Germany, the Netherlands, and Brazil, among others. All included studies reported miRNA expression profiles. Quantitative real-time polymerase chain reaction (qRT-PCR) was the primary detection method, while some studies utilized microarray or high-throughput sequencing techniques. Sample sources were diverse: human studies primarily used plasma and serum, whereas animal studies additionally included cardiac tissue samples. Detailed characteristics are presented in Table 1 and Table 2.

**Table 1.**
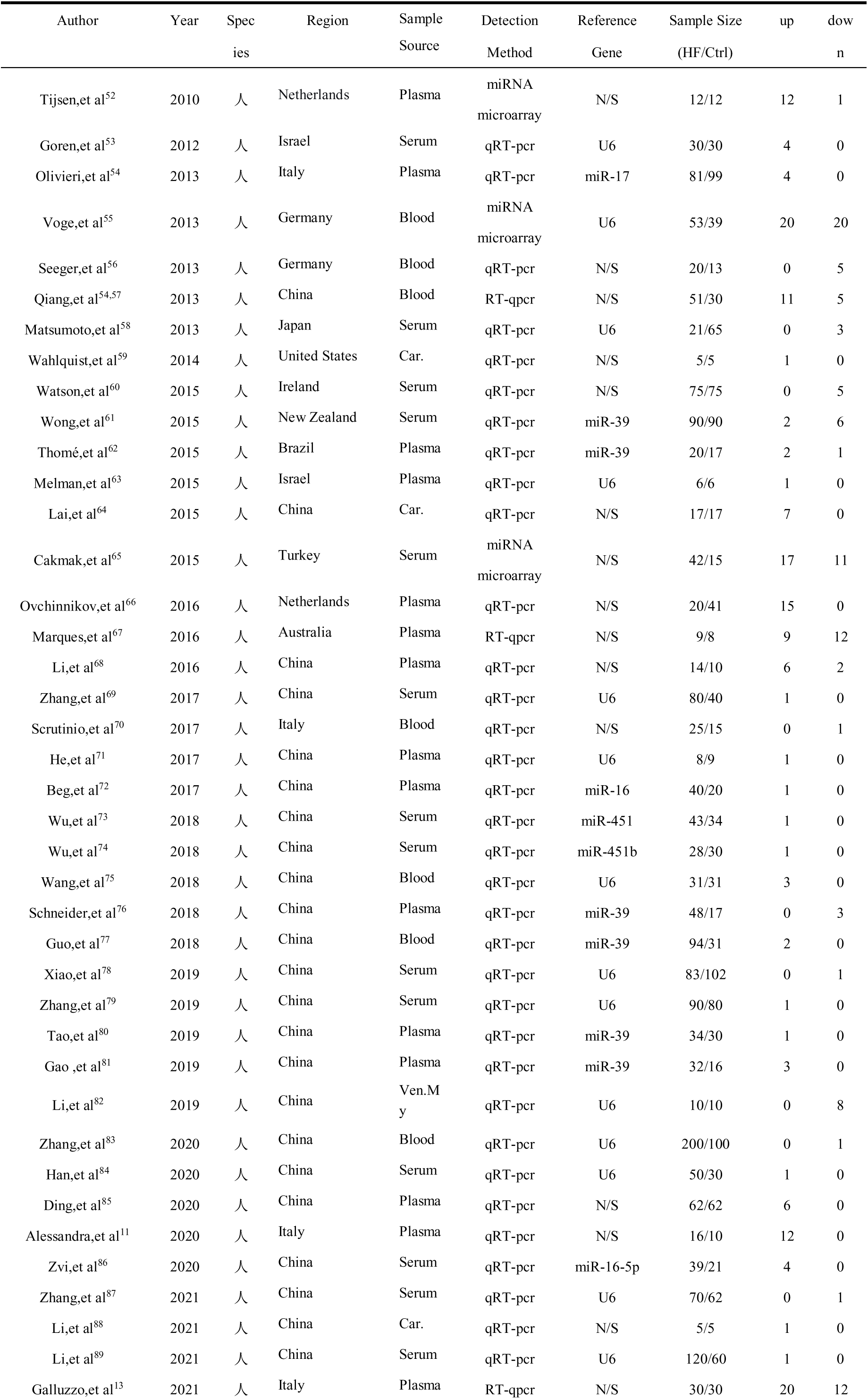
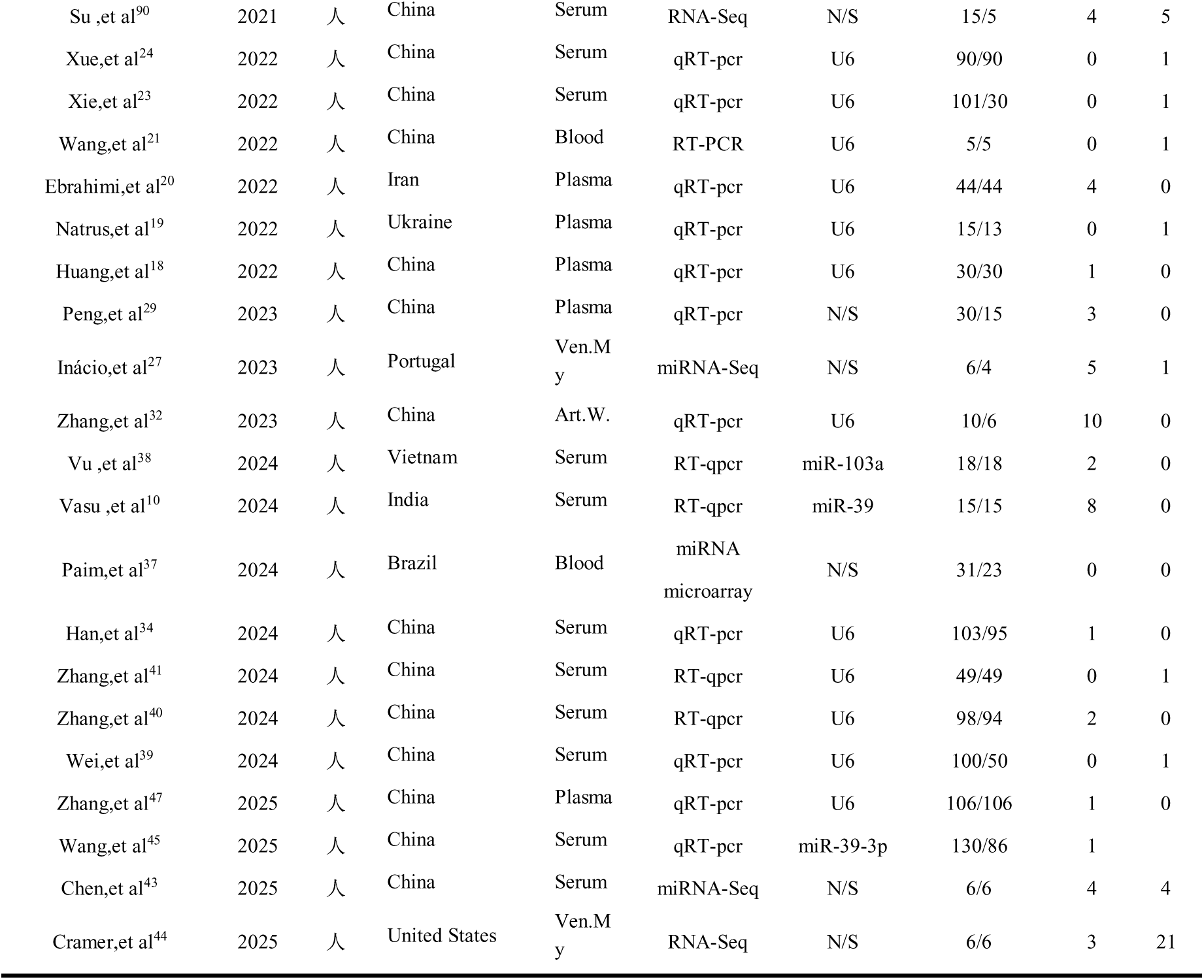
Characteristics of the included human studies on miRNA expression.

**Table 2.** Characteristics of the included animal studies on miRNA expression.

| Author | Year | Speci<br>es | Region | Sample<br>Source | Detection<br>Method | Reference<br>Gene | Sample Size<br>(HF/Ctrl) | up | down |
| --- | --- | --- | --- | --- | --- | --- | --- | --- | --- |
| B.Dickinson,et al<sup>91</sup> | 2013 | rats | USA | Plasma | qRT-pcr | U6 | 10/10 | 6 | 0 |
| K.Endo,et al<sup>92</sup> | 2013 | rats | Japan | Plasma | RT-qpcr | U6 | 13/9 | 1 | 0 |
| J.Zhou,et al<sup>93</sup> | 2015 | rats | China | Ven.My | RT-qpcr | U6 | 5/5 | 1 | 0 |
| H.Sang,et al<sup>94</sup> | 2015 | rats | China | Car. | qRT-pcr | N/S | 6/6 | 0 | 1 |
| X.Liu,et al<sup>95</sup> | 2016 | rats | China | Car. | qRT-pcr | N/S | 35/15 | 3 | 1 |
| X.Wang,et al<sup>96</sup> | 2017 | mice | China | Ven.My | qRT-pcr | U6 | 12/12 | 21 | 8 |
| Y.Du,et al<sup>97</sup> | 2017 | rats | China | Car. | qRT-pcr | U6 | 5/5 | 6 | 1 |
| L.Wong,et al<sup>98</sup> | 2017 | sheep | New<br>Zealand | Serum | qRT-pcr | N/S | 6/13 | 10 | 0 |
| S.Jung,et al<sup>99</sup> | 2018 | dogs | USA | Plasma | qRT-pcr | N/S | 8/9 | 4 | 4 |
| C.Tian,et al<sup>100</sup> | 2018 | rats | Netherla<br>nds | Car. | TaqMan<br>micRNA | U6 | 6/6 | 3 | 0 |
| Q.Su,et al<sup>101</sup> | 2019 | rats | China | Ven.My | qRT-pcr | U6 | 8/8 | 0 | 1 |
| G.Wang,et al<sup>102</sup> | 2020 | rats | China | Car. | qRT-pcr | U6 | 8/8 | 0 | 1 |
| Y.Yang,et al<sup>103</sup> | 2020 | rats | China | Plasma | qRT-pcr | U6 | 15/12 | 33 | 7 |
| W.Zhang,et al<sup>104</sup> | 2021 | mice | China | Car. | RT-qpcr | N/S | 6/6 | 0 | 1 |
| X.Zhou,et al<sup>25</sup> | 2022 | mice | China | Car. | miRNA-Seq | N/S | 5/5 | 1 | 1 |
| Y.Xiao<sup>22</sup> | 2022 | rats | China | Serum | qRT-pcr | miR-39-3p | 5/5 | 1 | 2 |
| Y.Wu,et al<sup>31</sup> | 2023 | rats | China | Ven.My | qRT-pcr | U6 | 6/6 | 1 | 0 |
| L.He,et al<sup>26</sup> | 2023 | mice | China | Ven.My | qRT-pcr | U6 | 6/6 | 1 | 0 |
| M.Ruppert,et al<sup>30</sup> | 2023 | rats | Hungary | Car. | miRNA-Seq | N/S | 5/8 | 44 | 6 |
| S.Mi ,et al<sup>28</sup> | 2023 | mice | China | Car. | qRT-pcr | U6 | 12/12 | 0 | 1 |
| Y.Zheng,et al<sup>42</sup> | 2024 | rats | China | Ven.My | qRT-pcr | U6 | 8/8 | 1 | 0 |
| Z.Gu,et al<sup>33</sup> | 2024 | rats | China | Car. | qRT-pcr | U6 | 8/8 | 0 | 1 |
| X.Liu,et al<sup>36</sup> | 2024 | mice | USA | Car. | qRT-pcr | U6 | 4/5 | 1 | 0 |
| Y.Hu ,et al<sup>35</sup> | 2024 | mice | China | Car. | qRT-pcr | N/S | 5/5 | 0 | 4 |
| J.Xu,et al<sup>46</sup> | 2025 | mice | China | Car. | RT-qpcr | N/S | 4/4 | 0 | 1 |

### Quality Assessment of Included Studies

The methodological quality of the included literature was assessed using standardized scales. For animal experimental studies, the SYRCLE risk of bias tool was applied. The results indicated that all studies performed well in key methodological domains such as sequence generation, allocation concealment, and completeness of outcome data, with most items rated as “Y” (Yes), reflecting an overall low risk of bias. For human case–control studies, a modified version of the Newcastle–Ottawa Scale was used. The majority of studies scored above 6 points (out of 9), with several achieving 8 points or even the maximum score, demonstrating high quality in the selection of cases and controls, comparability, and exposure assessment. Detailed results are provided in Supplementary Tables 1 and 2.

In summary, the overall methodological quality of the included animal and clinical studies is reasonably sound, providing robust evidence to support subsequent analyses.

### Overall Analysis of Differentially Expressed miRNAs

Meta-analysis results identified 71 circulating miRNAs with consistent expression patterns in heart failure out of the 225 differentially expressed miRNAs analyzed. Among these, 58 were significantly upregulated and 13 were downregulated (see Supplementary Figures 1-71). Notably, miR-21 was the most significantly upregulated miRNA (logOR = 8.15, 95% CI: 7.55–8.74, P < 0.001). It was detected as significantly elevated in 15 independent studies, with a total sample size of 722, demonstrating remarkable stability and reproducibility. Furthermore, other miRNAs including miR-423-5p and miR-210 were also consistently upregulated across multiple studies, suggesting their potential as diagnostic biomarkers (Figure 2).

**Figure 2.**
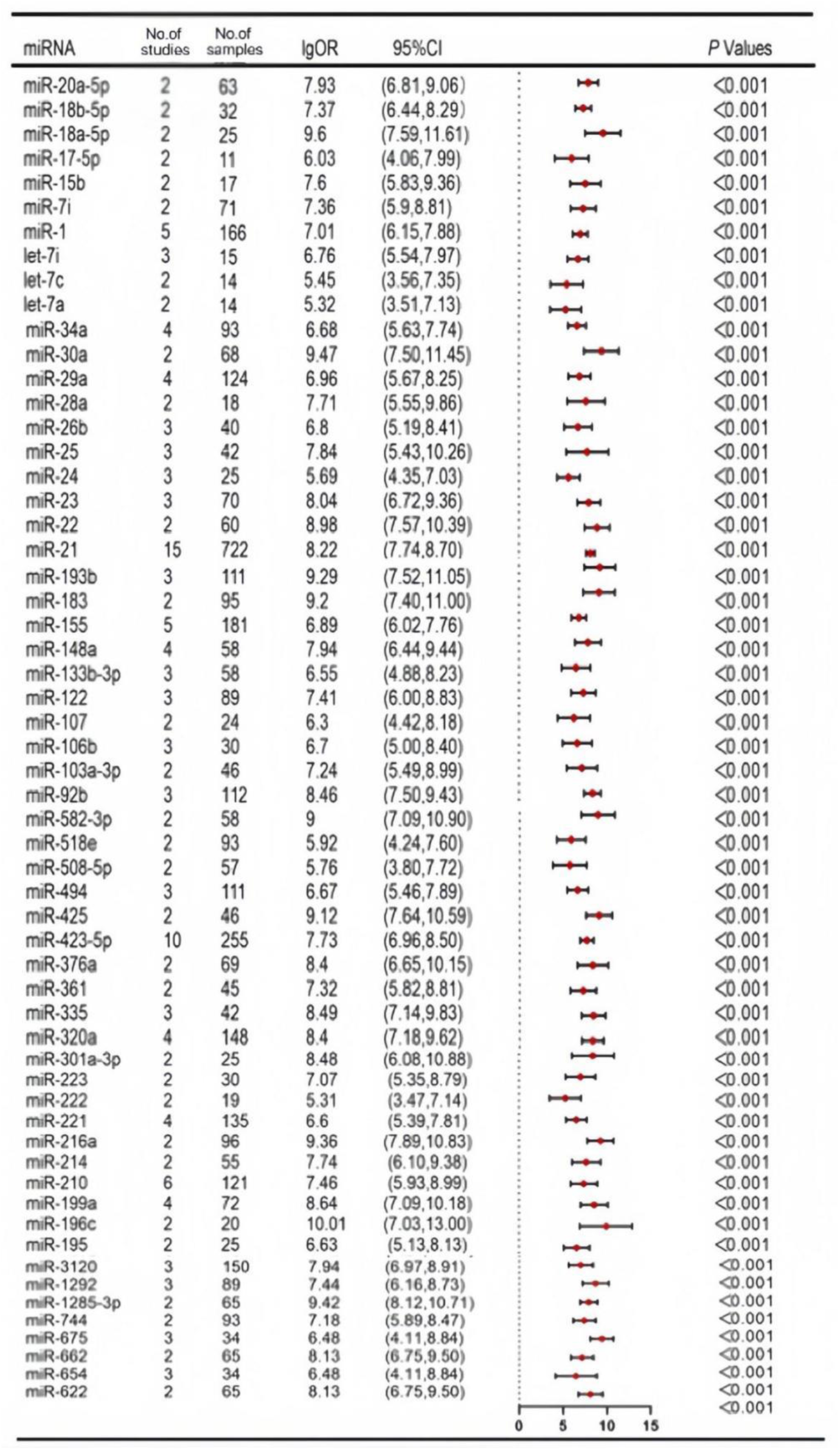
Statistically significant upregulated miRNAs.

Among the downregulated miRNAs (Figure 3), miR-144 showed the most pronounced alteration (logOR = -6.62, 95% CI: -10.20 to -3.04, P < 0.001), demonstrating a significant downregulation trend based on four independent studies involving a total of 83 participants. Other miRNAs, including miR-129, miR-126, and miR-27a, also demonstrated consistent downregulation(Figure 3).

**Figure 3.**
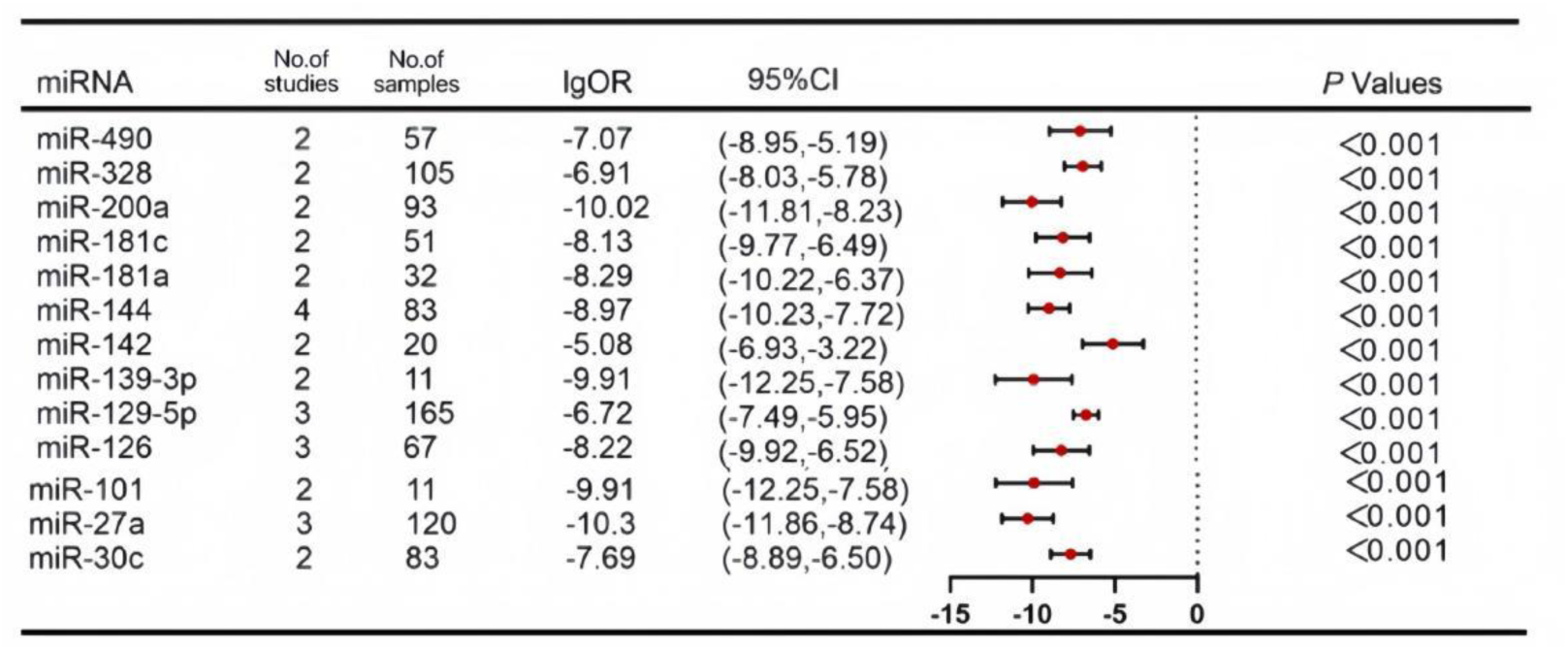
Statistically significant downregulated miRNAs.

### Subgroup Analysis

Across the 61 human studies, 45 aberrantly expressed miRNAs were identified, of which 36 were upregulated and 9 were downregulated (Supplementary Table 3). MiR-21 demonstrated the most pronounced upregulation. MiR-144 was consistently detected as downregulated in multiple studies. Regarding animal studies, 15 significantly upregulated miRNAs were identified, with no downregulated miRNAs observed (Supplementary Table 4). Among these, miRNAs including miR-106b, miR-199a, miR-210, miR-29a, and miR-196c were consistently upregulated across two or more studies with substantial effect sizes, suggesting their potential role in pathological regulation within animal models.

Based on the geographical origin of the study populations, 41 studies were from Asian countries and 20 from non-Asian countries. In the Asian subgroup, 29 differentially expressed miRNAs were identified (21 upregulated, 8 downregulated; Supplementary Table 5). MiR-21 showed the most significant upregulation, while miR-126 and miR-144 were consistently downregulated across multiple studies. In the non-Asian subgroup, 15 differentially expressed miRNAs were identified (14 upregulated, 1 downregulated; Supplementary Table 6). MiR-423-5p was reported in the highest number of studies, and miR-181c was the sole downregulated miRNA. Notably, miR-21, miR-148a, miR-221, miR-423-5p, and miR-622 exhibited consistent upregulation in both Asian and non-Asian populations, indicating cross-ethnic stability.

Based on sample source, 7 studies used tissue samples (myocardium, heart, or left ventricle), and 54 studies used blood samples (including serum, plasma, or peripheral blood), specifically 25 serum and 20 plasma studies. In human tissue samples, only 2 upregulated miRNAs (miR-25 and miR-148a) were identified, with no downregulated miRNAs observed (Supplementary Table 7). In human blood samples, 41 aberrantly expressed miRNAs were identified (34 upregulated, 7 downregulated; Supplementary Table 8). Regarding the magnitude of expression, miR-193b showed the most significant upregulation, while miR-21 was reported in the highest number of studies; miR-144 and miR-126 were consistently downregulated in multiple studies. Further analysis of plasma samples identified 13 aberrantly expressed miRNAs (12 upregulated, 1 downregulated; Supplementary Table 9). MiR-423-5p was the most significantly upregulated, and miR-126 was the sole downregulated miRNA. In serum samples, 5 aberrantly expressed miRNAs were identified (4 upregulated, 1 downregulated; Supplementary Table 10). MiR-21 showed the most significant upregulation, and miR-129-5p was the sole downregulated miRNA. Notably, miR-21 demonstrated consistent upregulation in both serum and plasma samples, further supporting its stability as a potential biomarker for heart failure.

### Evidence Quality Grading (GRADE) for Differentially Expressed miRNAs

In this study, the GRADE (Grading of Recommendations Assessment, Development, and Evaluation) system was employed to systematically and rigorously grade the quality of evidence for the key differentially expressed miRNAs identified through screening. This grading framework aims to evaluate the level of confidence in the estimated effect of each miRNA as a specific biomarker. The initial quality of evidence was pre-defined according to study design: all evidence derived from human observational studies (primarily case–control studies) started as “low” quality, whereas evidence from animal experimental studies was initially rated as “very low” quality due to inherent biological and clinical differences (i.e., indirectness) between the model systems and human disease states.

Subsequently, the body of evidence for each miRNA was systematically evaluated based on five predefined domains that could lower the quality of evidence(Supplementary Table 11): risk of bias, inconsistency, imprecision, indirectness, and publication bias. Additionally, three factors that could raise the evidence level were considered: large effect size, consistency across studies or subgroups (cross-validation), and stable expression across different sample types (stability). Ultimately, the evidence for each miRNA was comprehensively classified into one of four distinct quality grades: High, Moderate, Low, or Very Low.

Specifically, the GRADE assessment results revealed a clear distribution pattern (Figure 4).

**Figure 4.**
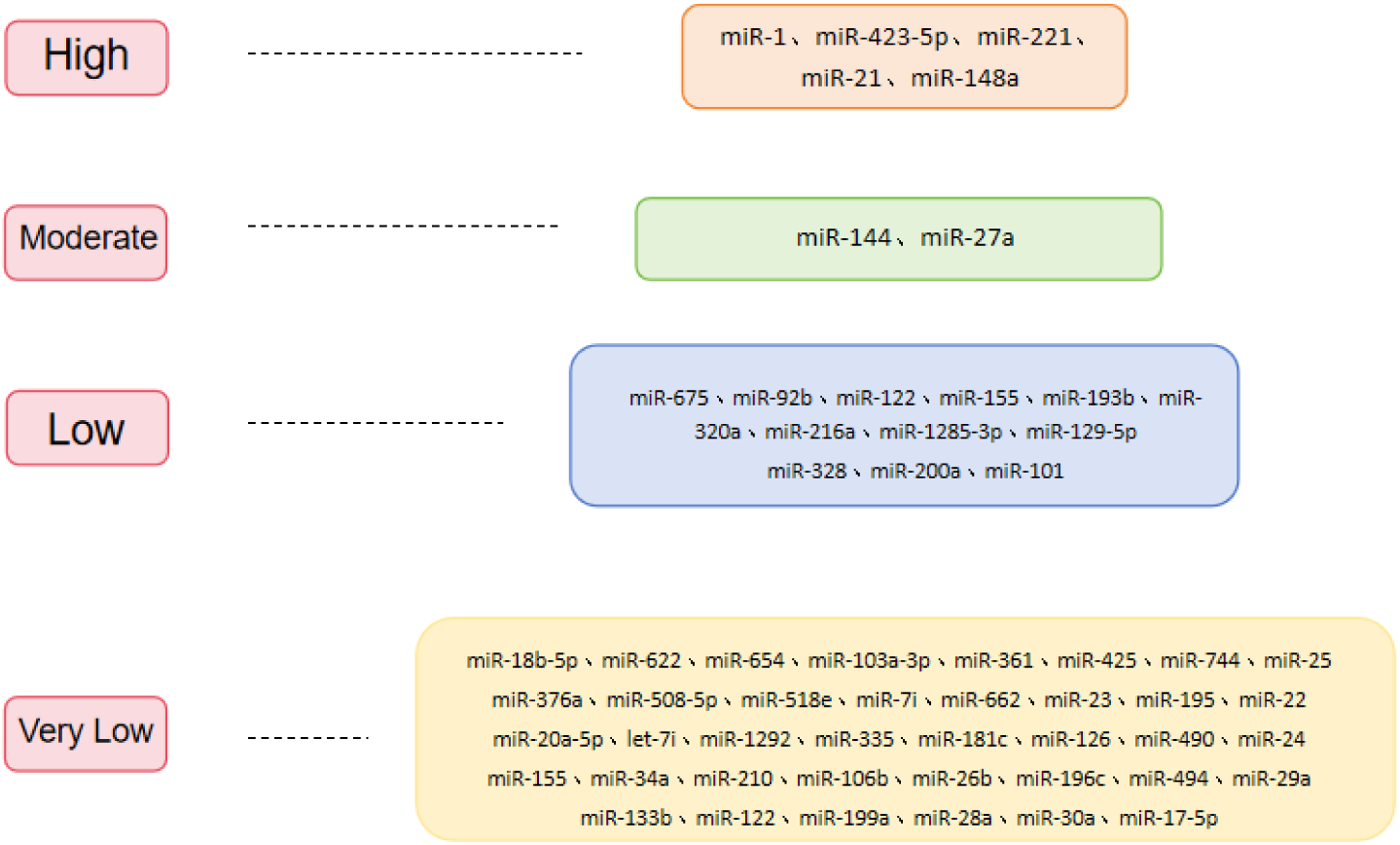
Hierarchical Assessment of miRNA Biomarkers for Heart Failure.

High-quality evidence: A total of 5 miRNAs—miR-1, miR-21, miR-221, miR-423-5p, and miR-148a—were rated as high quality. These represent the most robust and reliable core findings of this study. These miRNAs have been consistently replicated across multiple independent studies (e.g., miR-21 was supported by 13 studies with a total sample size of 701 cases) and generally have sufficient total sample sizes (typically >100), thereby avoiding downgrading for imprecision. Furthermore, most of them met upgrading criteria such as “cross-group consistency” or “multi-study validation,” indicating that their dysregulated expression patterns are reproducible across different populations or studies. Consequently, we have high confidence in the estimated effects of these five miRNAs as potential biomarkers.

Moderate-quality evidence: miR-144 and miR-27a were graded as moderate quality. Although supported by a number of studies, these miRNAs did not meet the high-quality threshold due to relatively limited sample sizes or insufficient upgrading factors, resulting in moderate certainty of the evidence.

Low- and very low-quality evidence: miRNAs such as miR-92b, miR-155, and miR-320a were rated as low quality, mainly constrained by small sample sizes or lack of upgrading factors. The largest number of miRNAs fell into the very low-quality category, forming the base of the evidence pyramid. This group primarily consists of two types: first, miRNAs reported in human studies but supported by few studies with very small total sample sizes (typically <100 cases), such as miR-34a and miR-210, which were downgraded primarily for imprecision; second, all miRNAs derived from animal studies—including miR-24, miR-155 (animal), and miR-34a (animal)—were uniformly rated as very low quality. This is largely due to inherent limitations in “population/model indirectness” in animal research, combined with generally small sample sizes, leading to considerable uncertainty in the evidence.

The entire GRADE assessment process was conducted independently by two researchers, and any disagreements were resolved through discussion and consensus, ensuring objectivity and rigor. In summary, this grading successfully identified a small set of high-confidence priority candidate miRNAs (e.g., miR-1, miR-21, etc.), providing key and robust anchors for subsequent biomarker translation and application. Simultaneously, it clearly revealed substantial weaknesses in the current evidence base, highlighting specific directions for future research to improve evidence quality through expanded clinical validation and optimized experimental design.

### Sensitivity Analysis

The sensitivity analysis was conducted to assess the robustness of the findings, specifically focusing on the potential impact of small-sample studies on the overall conclusions. Using the one-study-removed method, we systematically excluded 11 studies with small sample sizes (n ≤ 10) and re-analyzed the remaining 75 studies. The results identified 68 statistically significant miRNAs, comprising 55 upregulated and 13 downregulated miRNAs. Notably, 3 miRNAs that did not reach significance in the initial overall analysis showed statistical significance in this sensitivity analysis. Upon comparing the sensitivity analysis results with the overall analysis, we found that the overall pattern of the miRNA expression profile remained stable, with only minor differences. This indicates that small-sample studies had a limited impact on the overall conclusions of this systematic review, and our primary findings regarding the HF-related miRNA expression profile are relatively stable. The sensitivity analysis further supports the reliability of this study’s conclusions.

## Discussion

This systematic review and meta-analysis integrating the most recent evidence to evaluate the diagnostic value of circulating miRNAs in heart failure. It reaffirms the stability of previously reported core miRNAs, identifies additional candidates, and applies multi-dimensional stratification to guide clinical prioritisation.

Compared with previous systematic reviews^9,17^ this analysis, incorporating an expanded dataset and contemporary studies, confirms the stability of multiple core miRNA expressions. miR-21 and miR-423-5p, for example, showed persistent upregulation across additional independent studies and larger cohorts. with pooled effect sizes (logOR) comparable to prior estimates.These observations reinforce their utility as reliable diagnostic markers. Simultaneously, the newly identified miRNAs, such as miR-101,contribute to the diagnostic landscape despite support from fewer studies; their biological functions warrant further exploration in relation to HF pathological processes.

Subgroup analyses revealed substantial heterogeneity in the HF-related miRNA expression profiles, with important implications for the precise application of future biomarkers. Firstly, interspecies differences were evident: human studies frequently identified downregulated miRNAs, whereas animal models predominantly showed upregulation. likely reflecting limitations in replicating human pathophysiology and underscoring the need for cautious extrapolation from preclinical data. Ethnic variations were also apparent; core miRNAs such as miR-21 and miR-423-5p were consistently upregulated in both Asian and non-Asian populations, demonstrating potential for cross-ethnic application, others like let-7i-5p and miR-129 were significantly dysregulated only in specific ethnicities, indicating the feasibility of developing ethnicity-specific diagnostic panels in the future. Finally, Sample source further modulated profiles: correlations between tissue and blood-derived data were modest, and even among blood samples, serum and plasma yielded divergent results. The consistent upregulation of miR-21 in both serum and plasma makes it a highly promising biomarker for translation, whereas other miRNAs’ significance varied by source. These findings emphasise the importance of standardised protocols and population-specific considerations in future applications.

This study employed the GRADE system to systematically grade the evidence quality of the identified differentially expressed miRNAs, aiming to provide a scientifically grounded priority order for clinical translation. The grading results revealed substantial variation in evidence strength across different miRNAs, primarily due to the combined influence of factors such as the number of supporting studies, sample size, consistency across populations/sample types, and study origin (human vs. animal).

It is noteworthy that some miRNAs showing significant differential expression in the overall analysis were assigned lower evidence grades in the GRADE assessment, such as miR-210. Although it exhibited a consistently upregulated trend across multiple studies, its evidence body was limited by a small sample size (only 3 human studies with a total of 87 samples) and the indirectness associated with animal-derived studies. Consequently, it was downgraded for imprecision and indirectness, ultimately being classified as very low quality. This finding suggests that while miR-210 may be biologically relevant to the pathological process of heart failure, the evidence for its clinical diagnostic value remains insufficient and requires further validation in larger, prospective human cohorts.

In contrast, miR-1, miR-221, and miR-148a, together with the confirmed miR-21 and miR-423-5p from this study, were rated as high-quality evidence. These miRNAs have not only been consistently replicated across multiple independent studies but also demonstrated consistency across ethnicities and sample types, fulfilling upgrading criteria such as cross-group consistency and multi-study validation.

Ultimately, candidate miRNAs were stratified into four evidence tiers. miRNAs with high certainty at the apex (including miR-21, miR-423-5p, miR-221, etc.), with the most substantial research support and cross-population stability, should be prioritized as core targets for clinical validation and standardization. The miRNAs with moderate certainty(e.g., miR-144、miR-27a) warrant follow-up in additional cohorts. while miRNAs with low and very low certainty represent exploratory targets.

This study has limitations. First, the search was restricted to English-language databases, which may introduce bias. Second, heterogeneity likely arose from differences in study design, control groups, detection methods, and analytical approaches. Although we employed a random-effects model and conducted sensitivity analyses, residual heterogeneity might still influence the results. Third, due to limited access to raw data, we were unable to perform more refined subgroup analyses based on HF subtype or aetiology.

Future studies should prioritise large-scale, prospective, multicentre validations of high-priority microRNAs; investigations of expression patterns across heart failure phenotypes and aetiologies; and efforts to standardise detection and integrate microRNAs into multi-biomarker strategies with established markers such as BNP/NT-proBNP.

## Conclusion

In summary, this systematic review and meta-analysis identifies a panel of core circulating miRNAs with stable expression in HF and screened promising novel candidate molecules. By establishing a priority stratification through a comprehensive evaluation system, it provides clear guidance for subsequent research resource allocation and clinical translation pathways. The heterogeneity in expression revealed by subgroup analyses delineates the path toward precision diagnostics. Although challenges in standardisation and validation persist, these findings establish an evidence base for advancing microRNA-based approaches in heart failure.

